# Post-translational Modification of α-Synuclein Modifies Monomer Dynamics and Aggregation Kinetics

**DOI:** 10.1101/2024.05.06.592473

**Authors:** Kasun Gamage, Binyou Wang, Eldon R Hard, Thong Van, Ana Galesic, George R Phillips, Matthew Pratt, Lisa J. Lapidus

**Affiliations:** Department of Physics and Astronomy, Michigan State University, East Lansing, MI 48824; Department of Chemistry, University of Southern California, Los Angeles, CA 90089

## Abstract

The intrinsically disordered protein α-Synuclein is identified as a major toxic aggregate in Parkinson’s as well as several other neurodegenerative diseases. Recent work on this protein has focused on the effects of posttranslational modifications on aggregation kinetics. Among these, O-GlcNAcylation of α-Synuclein has been observed to inhibit the aggregation propensity of the protein. Here we investigate the monomer dynamics of two O-GlcNAcylated α-Synucleins, α-Syn(gT72) and α-Syn(gS87) and correlate them with the aggregation kinetics. We find that, compared to the unmodified protein, glycosylation at T72 makes the protein less compact and more diffusive while glycosylation at S87 makes the protein more compact and less diffusive. Based on a model of the earliest steps in aggregation, we predict that T72 should aggregate slower than unmodified protein, which is confirmed by ThT fluorescence measurements. In contrast, S87 should aggregate faster, which is not mirrored in ThT kinetics of later fibril formation but does not rule out a higher rate of formation of small oligomers. Together, these results show that posttranslational modifications do not uniformly affect aggregation propensity.

## Introduction

α-Syn can be subjected to a variety of posttranslational modifications (PTMs) that have the potential to affect the protein’s aggregation properties.[1-3] One PTM of particular interest is an intracellular form of glycosylation termed O-GlcNAc, which is the addition of the monosaccharide *N*-acetylglucosamine to serine and threonine residues.[4, 5] The O-GlcNAc modification is added by the single enzyme O-GlcNAc transferase (OGT) and removed by another called O-GlcNAcase (OGA).[6] This dynamic nature of the PTM enables it to respond to cellular inputs, and abnormal levels of O-GlcNAc have been linked with several human diseases.[7, 8] For example, O-GlcNAc levels in Alzheimer diseased brains are decreased by 40-50% in comparison to age-matched controls.[9-11] Additionally, we and others have shown that O-GlcNAc modification of aggregation prone proteins like α-Syn and Tau can inhibit the kinetics of amyloid fibril formation.[12-16] These discoveries have contributed to the development of a range of OGA inhibitors aimed at elevating O-GlcNAc levels in the brain,[17] and several of them slow protein-aggregate formation and onset of symptoms in animal models of Alzheimer’s or Parkinson’s disease.[12, 18] Some of these molecules have progressed to clinical trials; however, the exact mechanisms by which O-GlcNAc slows fibril formation are still poorly understood.

α-Syn has been found to be O-GlcNAc modified at up to nine different sites in proteomics experiments carried out on mouse or human brain tissue. Additionally, analysis of O-GlcNAc stoichiometry in a mouse-model of Parkinson’s disease found that 20% of α-Syn bears one O-GlcNAc modification and this ratio can be increased to 35% upon OGA inhibition,[18] but the amounts at each of the potential nine sites is still unknown. We have exploited protein semi-synthesis to a small panel of α-Syn proteins with site specific O-GlcNAc modification at T72, T75, T81 or S87.[14-16] As mentioned above, we found that O-GlcNAc generally slows α-Syn fibril formation, with interesting differences based on the site of modification. Specifically, O-GlcNAc modifications within the central core of the NAC region of α-Syn at T72, T75, or T81 were more inhibitory than modification at S87. More recently, we demonstrated that O-GlcNAc at S87 forces the formation of an alternate α-Syn fibril-structure that gives very little seeded aggregation in neurons and *in vivo*, leading to dramatically reduced pathology.[19] Structural analysis of these fibers using cryo-EM suggests that this is likely a steric effect of the O-GlcNAc modification blocking interactions that otherwise form in the unmodified protein. Separate molecular dynamic simulations by the Diao lab suggest that O-GlcNAc at T72, T75, or T81 slow aggregation by statically blocking inter-molecular interactions in the formation of early oligomers along the path to α-Syn fibril formation.[20]

In this work we investigate the kinetics of monomeric α-Syn and its glycosylates, well before fibril formation. We hypothesized that O-GlcNAc might also be slowing aggregation kinetically, by altering the intra-molecular diffusion of α-Syn monomers. Prior work has shown that the α-Synuclein monomer dynamics slow under aggregating conditions and this effect correlates with temperature, pH, mutation and small molecule aggregation inhibitors [21-25]. Slow reconfiguration can provide ample time for the formation of bimolecular interactions and speed up aggregation while faster reconfiguration rates can increase the rate of escape from an encounter complex, slowing down the aggregation process. To quantify the reconfiguration rate, we investigated the intramolecular diffusion of the protein using the Trp-Cys contact quenching technique where the α-Syn is mutated with a Cys and a Trp at positions 69 and 94 respectively. We find different effects of glycosylation at two different locations in the sequence that either speed up or slow down intramolecular diffusion.

## Results

### Synthesis of modified proteins

Measurement of intramolecular diffusion requires an IDP sequence to have one Trp and one Cys within 40 residues in the sequence. We have previously shown that the α-Synuclein mutant A69C F94W yields monomer dynamics similar to other loops in the chain and does not change the aggregation kinetics [22]. Our syntheses of α-Syn69C94W bearing O-GlcNAc at either T72 or S87, respectively termed α-Syn(gT72) and α-Syn(gS87), were generally similar to those that we have already published.[16-19] Specifically, we took advantage of expressed protein ligation (EPL),[26, 27] which enables the construction of proteins from recombinant and synthetic fragments bearing protein thioesters and *N*-terminal cysteine residues. In our previous syntheses, we deconstructed α-Syn into three fragments: *N*- and *C*-terminal recombinant proteins and a central glycopeptide containing the O-GlcNAc modification. We chose ligation sites, and thus the position of the required cysteine residues, at positions that are natively alanine in the α-Syn sequence. This allowed us to use a final desulfurization step to produce O-GlcNAc modified α-Syn with no primary sequence mutations. Here, we needed to alter these syntheses of the proteins to maintain the single C69 required for our biophysical analysis, and this necessitated a different approach for each protein.

In the case of α-Syn(gT72) (**Figure 1a**), our synthesis began with a recombinant protein thioester (**1**, residues 1-68), generated by fusion to an engineered intein from *Anabaena variabilis*, and a synthetic glycopeptide (**2**, residues 69-75) with a C-terminal hydrazide. We subjected these two fragments to ligation conditions, yielding intermediate **3** with C69 in place. We then generated protein **4** by protecting C69 using maleimide chemistry, preventing its loss during the final desulfurization step. Specifically, we used an *N*-alkyl-modified maleimide bearing six lysine residues, which allows for the separation of the protected product form any starting material using reverse-phase HPLC (RP-HPLC). We transformed the C-terminal hydrazide of **4** to a thioester using Dawson’s pyrazole method[28] and ligated it to recombinant fragment **5** (residues 76-140 containing W94) to give the full-length protein **6**. We subsequently converted C76 to α-Syn’s native alanine using radical-based desulfurization. Unfortunately, we found this reaction to be very sluggish compared to our previous syntheses and attribute this to the presence of the maleimide as the only differentiating feature. This slower reaction rate compromised the yield, and fairly notable amounts (> 3 mg) of protein are needed for the biophysical measurement. Therefore, we needed to use large amounts of **6** (15 mg) to give enough product. While this required us to perform the synthetic route several times, we were able to obtain the required quantities (3 mg) of α-Syn(gT72) after final removal of the maleimide group.

**Figure 1.**
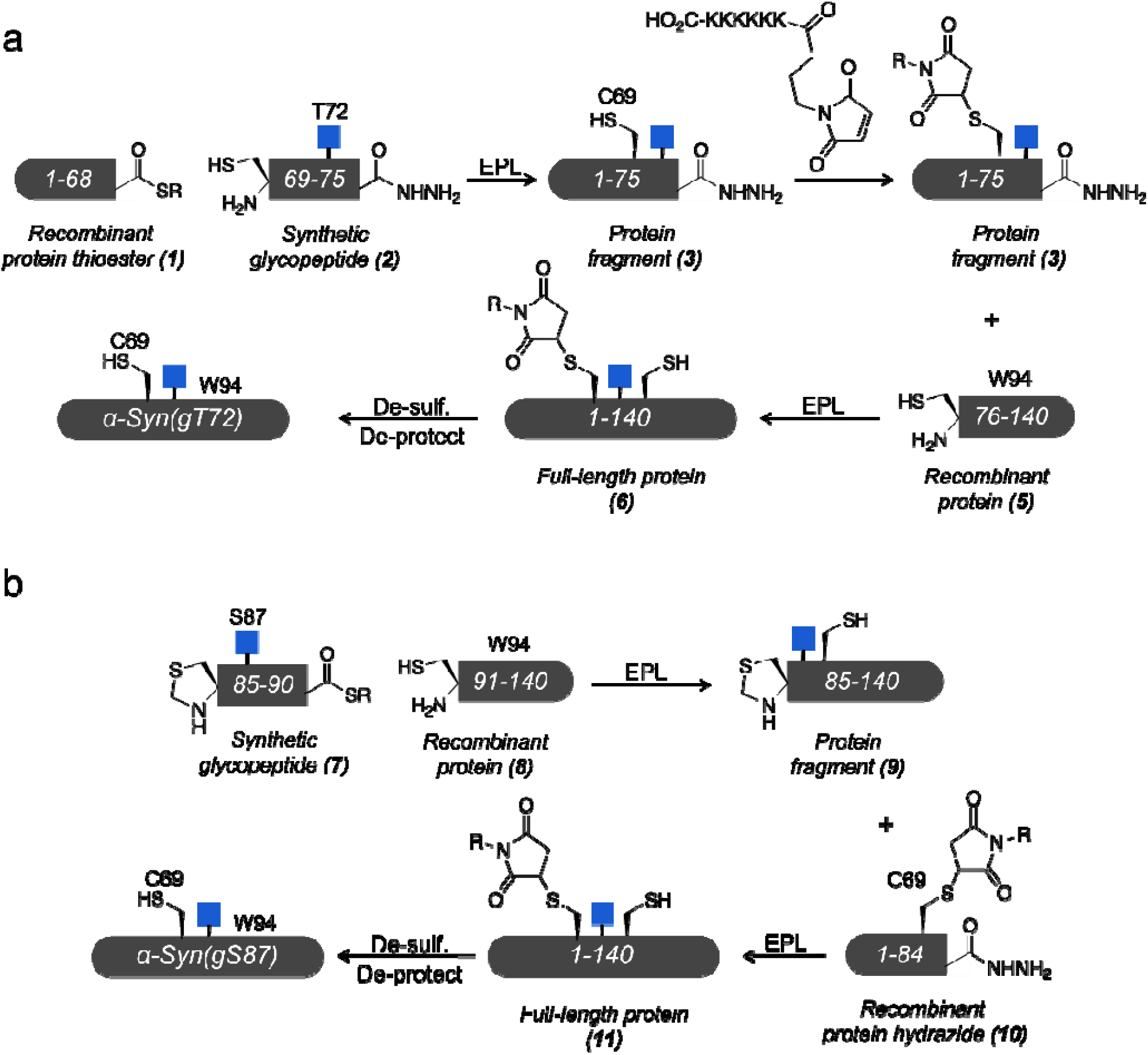
O-GlcNAc modified α-synuclein. a) Synthesis of α-Syn(gT72) with C69 and W94 using expressed protein ligation. b) Synthesis of α-Syn(gS87) with C69 and W94 using expressed protein ligation.

We took a different approach to α-Syn(gS87) (**Figure 1b**) and started with a synthetic glycopeptide thioester **7** (residues 85-90) and ligated it with the C-terminal recombinant fragment **8** (residues 91-140) to give intermediate **9**. In parallel, we prepared an *N*-terminal recombinant hydrazide (**10**, residues 1-84) containing C69 protected with the lysine-modified maleimide described above. We then deprotected the *N*-terminal thiazolidine of **9** and ligated it to **10** to generate the full-length intermediate **11**. We used the same desulfurization chemistry to transform C85 and C91 to the native alanine residues, but again found this action to be kinetically slow and low yielding. Again, this required us to go through this reaction sequence several times to give 25 mg of **11**. However, we were able to prepare the material and remove the maleimide protecting-group to give 9 mg of α-Syn(gS87). In both cases, we characterized the purify and identity of the proteins using RP-HPLC and ESI-MS on a high-resolution QTOF instrument (**Figur S1**).

### Aggregation Kinetics

Before moving to our analysis of the diffusion of these proteins, we first wanted to ensure that the inclusion of the necessary C69 and W94 mutations did not notably affect the relative fibrilization kinetics of these O-GlcNAc modified proteins compared to each other and unmodified α-Syn. Accordingly, we subjected these three proteins at 50 μM concentration in phosphate buffered silane (PBS) to aggregation conditions (agitation at 37 °C) in a plate reader. We also added the amyloid-sensitive dye Thioflavin T (ThT, 10 μM) and measured fluorescence every 15 min over 7 days (**Figure 2**). The results were highly consistent with our previous analysis [16]. We observed very little if any aggregation by α-Syn(gT72) over the course of the assay, while the kinetics of α-Syn(gS87) fibril formation is delayed compared to unmodified protein. We also calculated the onset times for aggregation (2.5 times the initial ThT value) for α-Syn and α-Syn(gS87) and confirmed that the differences in aggregation kinetics were statistically significant.

**Figure 2.**
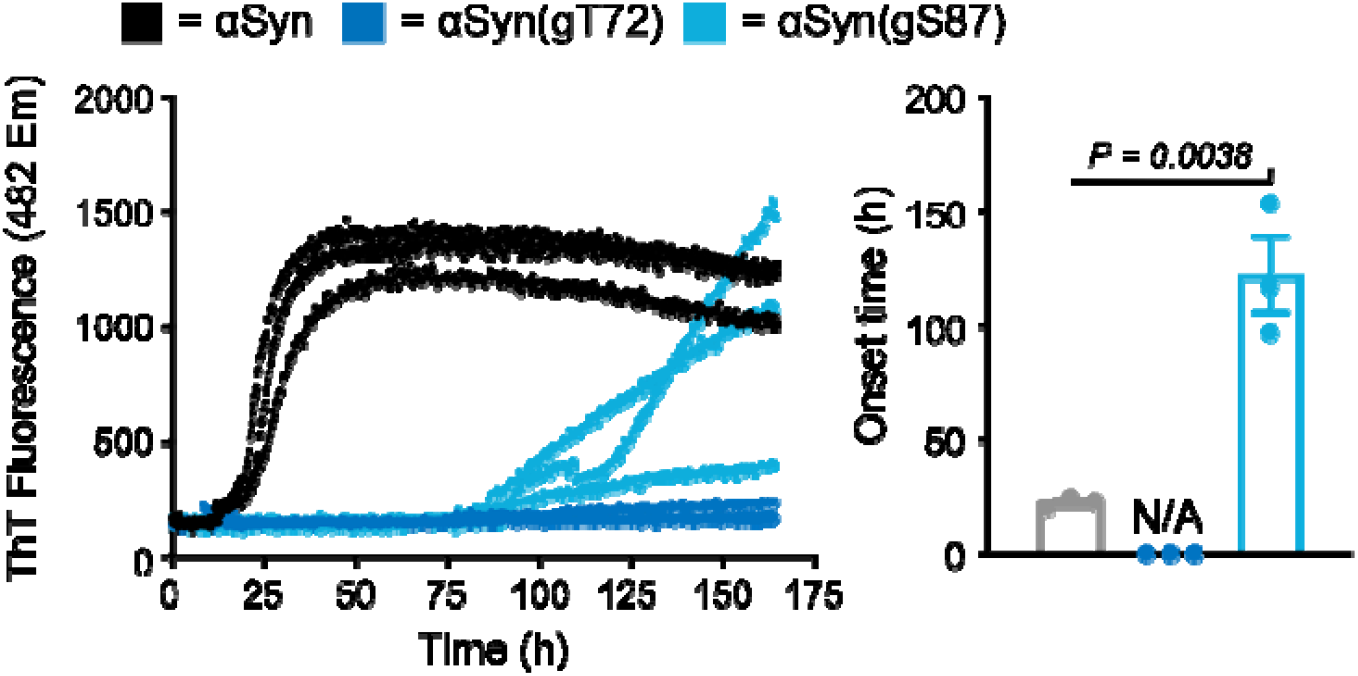
Kinetics of α-Syn fibril formation. The indicated α-Syn proteins bearing C69 and W94 (50 μM) where subjected to aggregation conditions and analysis using ThT fluorescence (λ_ex_ = 450 nm, λ_em_ = 482 nm). b) The fibrillization onset-times were calculated by measuring the time required for fluorescence to reach 2.5-times the initial reading. Onset-time results are mean ±SEM of experimental replicates (n=3). Statistical significance was determined using a two-way, unpaired Student’s t-test.

### Monomer Dynamics

To investigate the intramolecular diffusion of the proteins, the Trp is excited to a long-lived triplet state which then can be quenched by the Cys at close contact using a 2-laser, pump-probe instrument (Figure S2). The observed decay of the Trp is kinetically modeled (**Figure 3**), where the two contacts can diffuse towards each other at a rate of *k*_*D+*_ (diffusion-limited rate) and quench at a rate of *q* or diffuse away from each other at a rate of *k*_*D-*_. The observed decay rate is given by[21]

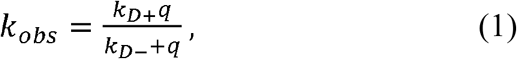

which can be rewritten as

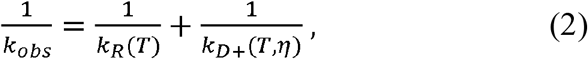

where

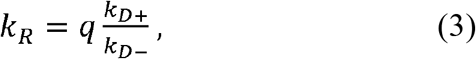

is the reaction-limited rate. Here we assume that *k*_*R*_ depends only on temperature while *k*_*D+*_ depends on temperature as well as viscosity η. Therefore, by measuring *k*_*obs*_ for different η at a particular temperature, a linear relationship is obtained between *1/k*_*obs*_ and η where *1/k*_*R*_ and *1/k*_*D+*_ can be determined from the intercept and the slope (normalized by the viscosity at 37°C).[29]

**Figure 3.**
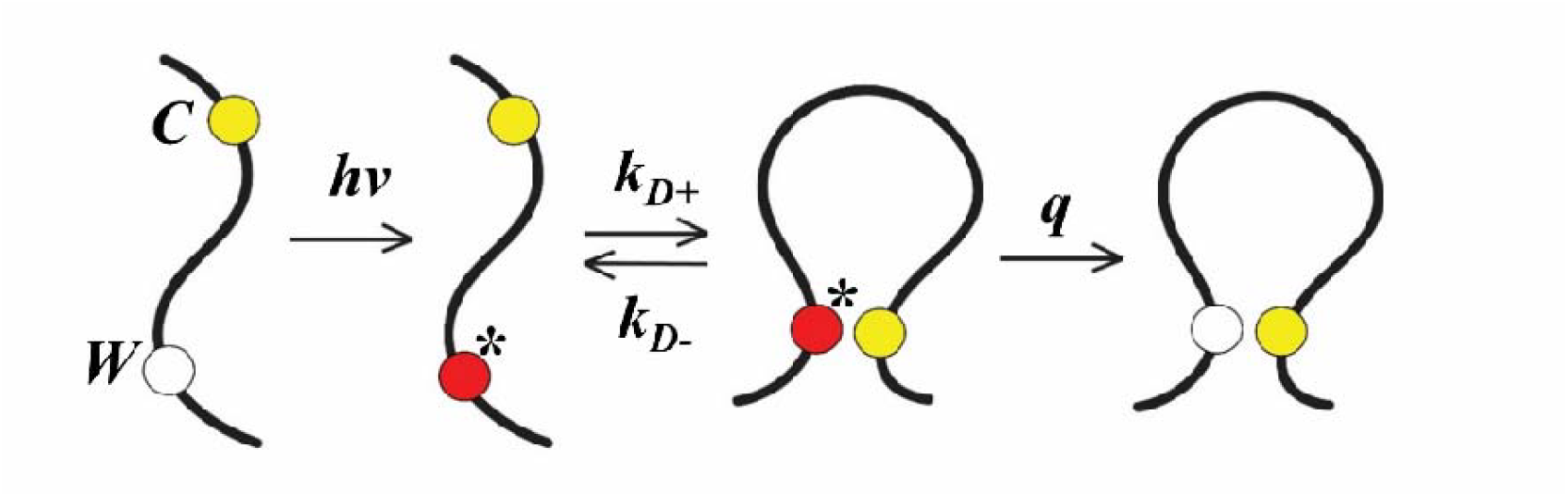
Schematic depicting the Trp (*W*)-Cys (*C*) contact quenching model. UV radiation excites *W* to a triplet state where it diffuses towards *C* at a rate of *k*_*D+*_. At close contact *W* is either quenched at a rate of *q* or diffuse away at a rate of *k*_*D-*_. Excitation is marked as **\***.

The decay of Trp was recorded for the unmodified and modified proteins at 37 °C with various concentrations of sucrose and fit to 1^st^ order decays. **Figure 4** shows the *1/k*_*obs*_ versus the viscosity for each protein. The intercept of each line is equal to *1/k*_*R*_ and the slope of each line is equal to 1/*ηk*_*D+*_. For α-Syn(gS87) the intercept (1/*k*_*R*_ = -6.9x10^−9^ ± 1.5x10^−7^ s) is consistent with zero so we can only find a lower limit in *k*_*R*_. Similarly, the slope (*1/ηk*_*D+*_ = 9.8x10^−8^ ± 1.7x10^−7^ s cP^-1^) of α-Syn(gT72) is also consistent with zero. Therefore, we assign the inverse of the maximum 95% confidence as the lower limit for both *k*_*R*_ of α-Syn(gS87) and *k*_*D+*_ of α-Syn(gT72). The diffusion-limited and the reaction-limited rates were extracted from the linear fits and plotted in **Figure 5(a) and 5(b)** respectively.

**Figure 4.**
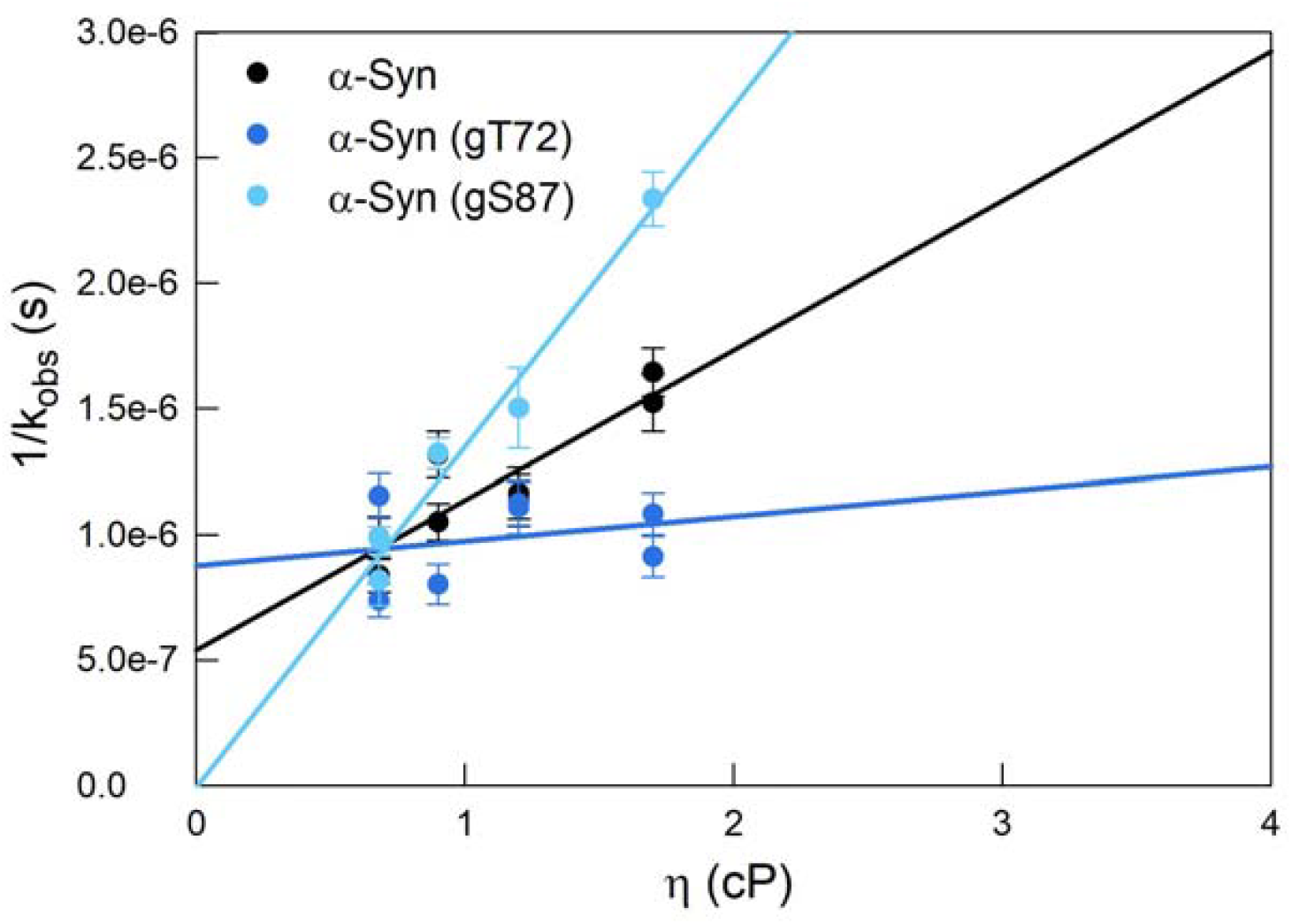
Observed decay rates. *1/k*_*obs*_ plotted against viscosity for α-Syn, α-Syn(gT72) and α-Syn(gS87) at 37□. Rates and their standard errors were obtained from 1^st^ order decay fits.

**Figure 5.**
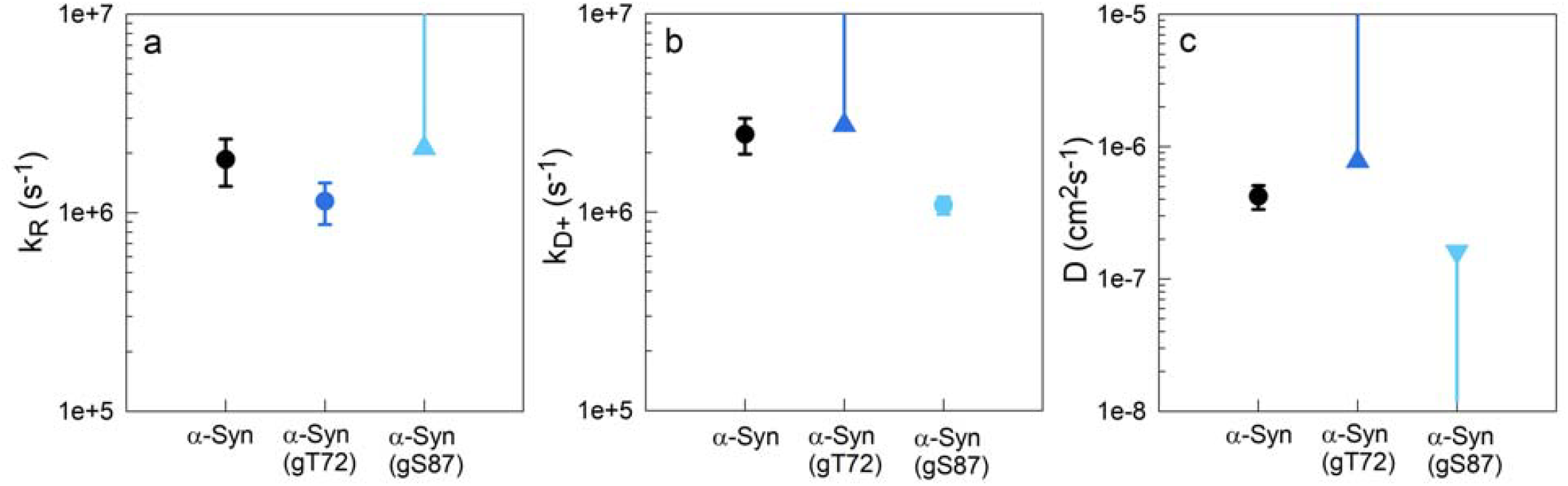
Computed rates and diffusion coefficients for α-Syn, α-Syn(gT72) and α-Syn(gS87) at 37°C. a) Reaction-limited rates. b) Diffusion-limited rates calculated for η = 0.68cP (37°C in water). c) Diffusion coefficients. Triangles indicate the lower or upper limits computed in cases where the 1/*k*_*R*_ and 1/*k*_*D+*_ values were consistent with zero.

To calculate the intramolecular diffusion coefficient from these measured rates we follow the SSS theory.[30] The theory considers a 1-D potential governed by the distance probability distribution *P(r)* between the Trp and the Cys. The reaction-limited and diffusion-limited rates are given by

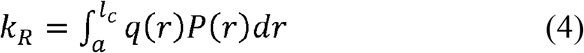

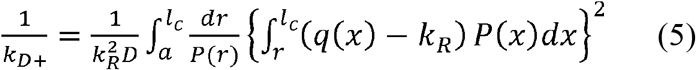

where *a* = 4Å is the van der Waals contact distance, *l*_*c*_ is the contour length of the chain, *r* is the distance between the Trp and the Cys, the distance dependent quenching rate *q*(*r*) = 4.2 x 10^9^ exp(4.0(*r*-*a*)) *s*^-1^ determined experimentally [31] and D is the diffusion coefficient. Here we assume a Gaussian chain model with a normalized probability distribution of,

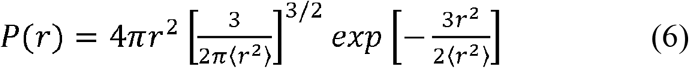

where ⟨*r*^2^ ⟩is the mean squared contact distance.

Using Eqs. 4 and 6, we can find an average distance between C69 and W94 that yields a *k*_*R*_ that agrees with the measured values. We find that *<r*^*2*^_α*-Syn(gT72)*_*>* is greater than *<r*^*2*^_*α-Syn*_*>*, which is greater than *<r*^*2*^_*α-Syn(gS87)*_*>*. Eq. 6 is then employed to calculate the intramolecular diffusion coefficient, *D*, using the measured *k*_*D+*_, as shown in **Figure 5(c)** and **Table S2**. Due to the measurement limitations described above, we estimate the lower limit of *α*-Syn(gT72) and an upper limit of *α*-Syn(gS87). Overall we find *D*_*α-Syn(gT72)*_ > *D*_*α-Syn*_ > *D*_*α-Syn(gS87)*_.

## Discussion

We see significant differences in diffusion among the proteins at their monomeric state. *α*-Syn(gT72) diffuses faster compared to *α*-Syn(gS87) and the unmodified *α*-Syn making it less prone to aggregation. This resistance to aggregation is maintained even after several days as observed through ThT fluorescence (**Figure 2**). On the other hand, *α*-Syn(gS87) reconfigures slower than the unmodified *α*-Syn which could make it more prone to aggregation in its monomeric state, but this behavior is not mirrored at longer times (**Figure 2**) where *α*-Syn(gS87) exhibits a significant delay in forming fibrils compared to the unmodified *α*-Syn. However, ThT fluorescence is insensitive to small oligomer formation and this modification may inhibit the oligomer-to-fibril transition. Indeed, previous work has found that the fibrils of *α*-Syn(gS87) are distinct from unmodified fibrils.[19]

To understand the implications of the measured diffusion coefficients for unmodified and modified *α*-Synuclein, we employ a kinetic model of early aggregation, shown in **Figure 6**. The model assumes the conformations of monomeric *α*-Synuclein can be divided into two populations, non-aggregation-prone (M) and aggregation-prone (M*), which interconvert at rate proportional to *D*. An aggregation-prone conformation likely has more solvent-exposed hydrophobic residues than a non-aggregation prone conformation. When two aggregation-prone monomers meet due to bimolecular diffusion (at rate *k*_*bi*_), they can form an encounter complex that can either go on to form stabilizing interactions in an oligomer (O) or disassociate due to one M* converting to M with rate *k*_*-1*_. Thus, the rate of oligomer formation depends on the relative values of *k*_*-1*_ and *k*_*bi*_, assuming these rates are much faster than *k*_*olig*_. Subsequent steps of fibril nucleation and elongation may differ between modified and unmodified proteins but are ultimately kinetically controlled by the faster rates.

**Figure 6.**
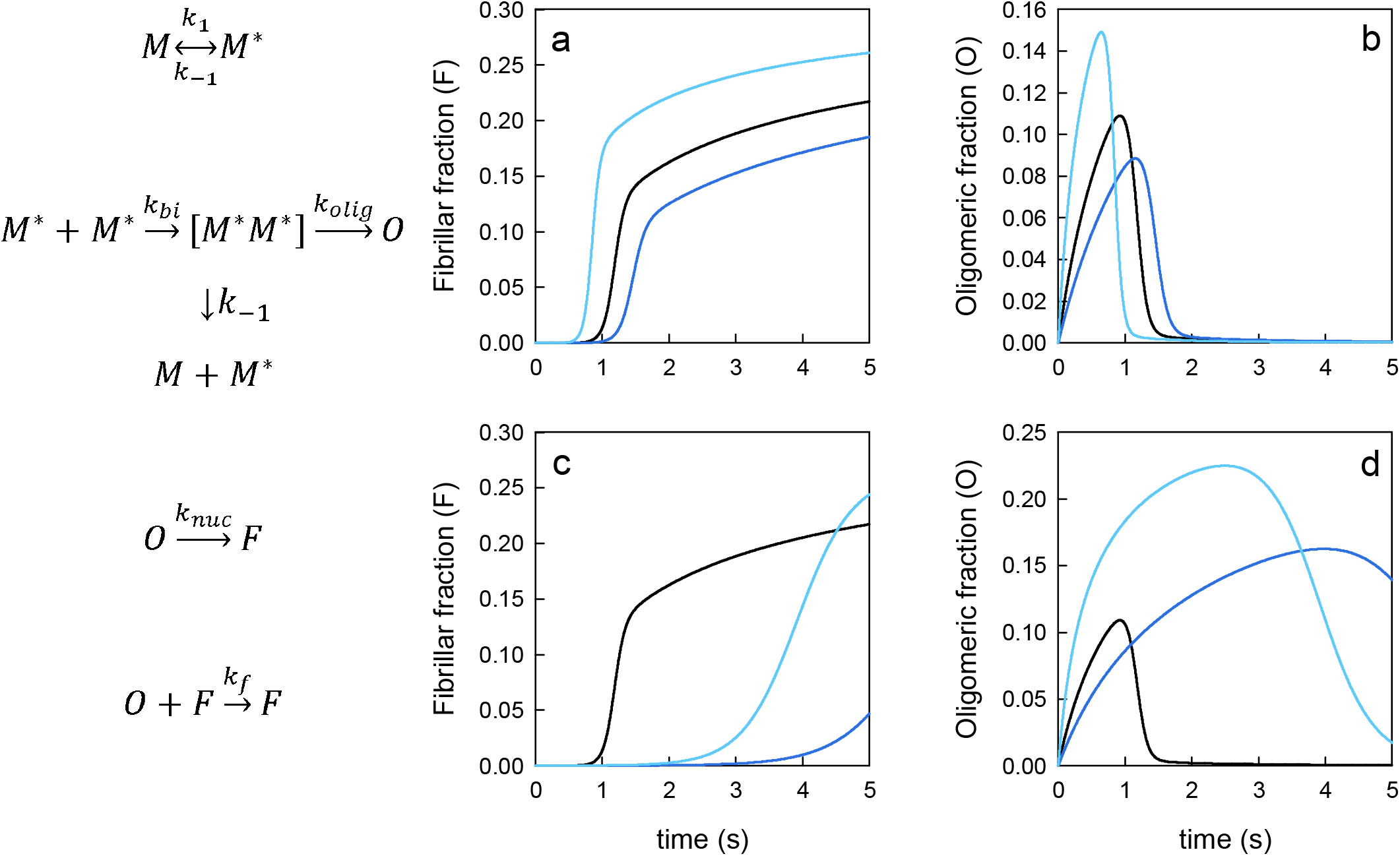
Kinetic model of aggregation. a) Formation of fibrils and b) formation of oligomers for *k*_*1*_*=k*_*-1*_ = 4.7x10^6^ s^-1^ (black, unmodified protein), *k*_*1*_*=k*_*-1*_ = 7.8x10^6^ s^-1^ (dark blue, a-syn(gT72)) and *k*_*1*_*=k*_*-1*_ = 1.6x10^6^ s^-1^ (light blue, a-syn(gS87)). All other rates are the same for each (*k*_*bi*_ = 9.7x10^4^ s^-1^, *k*_*olig*_ = 100 s^-1^, *k*_*nuc*_ = .001 s^-1^, *k*_*f*_ = 100 s^-1^). c) Formation of fibrils and d) formation of oligomers. The rates are same as for (a) and (b) except *k*_*f*_ = 10 s^-1^ for a-syn(gT72) and a-syn(gS87).

We can estimate the reconfiguration rates *k*_*1*_*=k*_*-1*_*=4D/(2R*_*G*_)^*2*^ as the time to diffuse across the diameter of the chain using the intramolecular diffusion coefficients shown in **Figure 5** and an estimated radius of gyration, *R*_*G*_ *∼* 3 nm [22], which should not change very much with modification. We can estimate *k*_*bi*_ = *4D/(2r*^*2*^*) =* 9.7 x 10^4^ s^-1^ where *r* is calculated from the concentration of the protein (50μM) and a typical intermolecular diffusion coefficient for a 14 kDa protein (*D*_*bi*_ = 1.0 x 10^−6^ cm^2^s^-1^). Solving this model for arbitrary rates of *k*_*olig*_ = 100 s^-1^, *k*_*nuc*_ = 0.001 s^-1^ and *k*_*f*_ = 100 s^-1^, the populations for the oligomer and fibril states are plotted in **Figure 6a and b**. *α*-Syn(gS87) has the fastest formation of O and F and *α*-Syn(gT72) has the slowest formation proportionate with the differences in *k*_*1*_. However, the fibril formation kinetics do not match the measured kinetics shown in **Figure 2**, indicating there is another difference in rates due to the PTM. **Figure 6c and d** show the fibril and oligomer formation if *k*_*f*_ is slowed to 10 s^-1^ for *α*-Syn(gT72) and *α*-Syn(gS87), which is qualitative agreement with **Figure 2**. The delay in fibril formation now extends the lifetime of the oligomers for the modified proteins.

The modeling results show that a PTM can either speed up or slow down the formation of oligomers, depending on the position and slow down the growth of fibrils, regardless of position. Cryo-EM data comparing unmodified and gS87 fibrils show significant structural differences that can be attributed to the presence of the glycosyl group. Therefore, fibril formation could be slowed simply by steric hindrance from the glycosyl group. This may explain why fibrils are delayed or eliminated for a variety of PTM locations in the range of residues 72-87 [16] because the variations observed in fibril formation may be entirely due to variations in intramolecular diffusion and consequent oligomer formation. However, the variable experimental results on monomer reconfiguration are less easily explained by steric hinderance because modification of T72 speeds up reconfiguration while modification of S87 slows it down. One possible explanation for the variability is that the most common intramolecular interactions for these two sites are different. If unmodified T72 makes many contacts far in sequence, then disruption of those contacts will make the chain more diffusive. Previous measurements of intramolecular diffusion on the *α*-Syn mutant T72P also showed an increase in intramolecular diffusion [32]. In contrast, if unmodified S87 makes more contacts nearby in sequence, then disruption of those contacts may allow other residues to make more contacts far in sequence, making the chain less diffusive. Molecular dynamics simulations of unmodified *α*-Synuclein support this hypothesis [33].

While it is clear that O-GlcNAc modification is overall neuroprotective [18, 34], it is not known whether all modified sites in *α*-Syn are equally populated or equally protective. The difference in monomer reconfiguration between gT72 and gS87 suggest there may be differences in protection, but it is still unclear which step of *α*-Syn aggregation is toxic to cells. There is much evidence that small oligomers are the toxic species in neurodegenerative diseases. However, the spread of fibrils in the brain by seeding is also important and gS87 fibrils have been shown to suppress seeding in neurons compared to unmodified fibrils [19]. Observing small oligomers is still extremely difficult in most experimental situations so knowledge of monomer aggregation propensity via reconfiguration dynamics is our best view of early aggregation processes.

## Methods

### General

All solvents and reagents were purchased from commercial sources (Sigma-Aldrich, VWR, GoldBio, EMD, P3 Bio Systems, etc.) and used without further purification. All aqueous solutions were prepared using ultrapure laboratory grade water (deionized, filtered, sterilized) obtained from an in-house Millipore water purification system. All growth media (LB broth, Miller) were prepared, sterilized, stored, and used according to the manufacturer. Stock solutions of antibiotics were made at a working concentration of 1000x (ampicillin sodium salt, Sigma Aldrich, 100 mg/mL, kanamycin sulfate, Carbosynth, 50 mg/mL) and stored at -20 °C. All bacterial growth media and cultures were handled under sterile conditions under open flame. Reverse phase high performance liquid chromatography (RP-HPLC) was performed using Agilent Technologies 1200 Series HPLC. The HPLC buffers used were 0.1% TFA in H2O (Buffer A), and 0.1% TFA, 90% ACN in H2O (Buffer B). Mass spectra were acquired on an Agilent LC QTOF MS/MS.

### Expression of recombinant full length *α*-Synuclein

BL21(DE3) chemically competent *E. coli* (VWR) were transformed with the pTXB1 construct containing wild-type human *α*-Synuclein, plated on LB agar plates containing 100μg/mL ampicillin (LB-amp), and incubated at 37 °C 16hrs. Single colonies were picked and used to inoculate two 50 mL LB-Amp cultures, which were grown at 37 °C with shaking at 225 rpm overnight. The 50mL cultures were combined and used to grow 500mL TB-Amp cultures. These cultures were grown to an OD600 of 0.6-0.8 at 37 °C shaking at 225 rpm, and then expression was induced with IPTG. (Isopropyl-ß-D-1-thiogalactopyranoside) (final concentration: 1mM) with shaking at 25 °C and 225 rpm for 16hrs. Bacteria were harvested by centrifugation (6,000 x g, 10 min, 4 °C), and the cell pellets were lysed by three freeze thaw cycles, using liquid N2 and a 37 °C water bath. Cell lysates were resuspended, in 10mL (per 500mL of culture) of lysis buffer (500mM NaCl, 100mM Tris, 10mM β-mercaptoethanol (βME), 2mM phenylmethanesulfonylfluoride (PMSF), 1mM EDTA, pH 8.0). Cell lysates were then tip sonicated at 70% amplitude for 5min 30s on/30s off, followed by clarification via centrifugation (20,000 x g, 30 min, 4 °C). The supernatant was acidified, on ice, to pH 3.5 with HCl and then incubated on ice for an additional 30 min before clarification again (20,000 x g, 30 min, 4 °C). The supernatant was collected and dialyzed against 3 x 1 L of 1% acetic acid in water (degassed with N_2_, 1 hr per L). The dialyzed protein solution was then purified by RP-HPLC over a C4 semi-preparative column (Phenomenex). Purified material was flash frozen in liquid N2 and lyophilized. Pure *α*-Synuclein was characterized by C4 analytical RP-HPLC column (Higgins Analytical) and ESI-MS (M+H+) and yield was determined by Pierce BCA assay (Thermo Scientific).

### Expression of *α*-Synuclein C-terminal fragment (5, 8)

BL21(DE3) competent *E. coli* (VWR) were transformed with the pET42b construct containing **5** or **8** by heat shock and plated on selective LB agar plates containing 50μg/mL kanamycin. Expression and purification of the fragments was carried out as described above for recombinant *α*-Synuclein.

### Expression of *α*-Synuclein N-terminal thioester/hydrazide (1, 10)

BL21(DE3) competent *E. coli* (VWR) were transformed with the modified pTXB1 construct containing **3** by heat shock, plated on LB agar plates containing 100μg/mL ampicillin, and incubated at 37 °C for 16 hr. Bacteria were cultured and induced as previously described. After harvesting bacteria by centrifugation (6,000 x g, 10 min, 4 °C), the cell pellet was resuspended on ice in 10 mL (per 500 mL of culture) in cold lysis buffer (50mM NaH2PO4, 300mM NaCl, 5mM imidazole, 2mM TCEP HCl, 2mM PMSF, pH 7.4) and lysed by tip sonication (70% amplitude, 30 sec pulse duration, 30 sec rest for 5 min) while on ice. The cell lysate was clarified by centrifugation (20,000 x g, 30 min, 4 °C) and the resulting supernatant was incubated with His-tag cobalt resin (GoldBio) for 1 hr. The column was washed with 20 column volumes (CV) of deionized water and wash buffer each prior to use. The column was washed with 15 x 1 CV of wash buffer 2 (lysis buffer, 50mM imidazole), and then eluted in 7 x 1.5 CV of elution buffer (lysis buffer, 250 mM imidazole). Elution fractions were dialyzed against 3 x 1 L (PBS (Cytiva HyClone), 1mM TCEP HCl, pH 7.2) using a 3.5 kDa cutoff (Amicon Ultra 3.5 kDa MW cut-off, Millipore). 2-Sodium mercaptoethane sulfonate (MESNa) was added to a final concentration of 200 mM along with fresh TCEP (2mM final concentration), and the thiolysis reaction was incubated at room temperature for 48 hrs to generate the protein thioester. For hydrazide **10**, 10% w/v hydrazine monohydrate was added, and the pH was lowered to 7 before allowing the reaction to proceed for 48hrs at room temperature. The thiolysis reaction was purified over a C4 semi-prep column and stored as a lyophilized solid. Pure thioester/hydrazide was characterized by analytical RP-HPLC and LC MS QTOF.

### Solid phase synthesis of hydrazide peptides 2 and 7

All solid-phase peptide syntheses were conducted manually using 2-chlorotrityl resin (P3 Bio Systems), with an estimated loading of 1.33mmol/g. The resins were functionalized with hydrazine by shaking the resin at 30 °C for 30min twice in 5% hydrazine monohydrate (Sigma Aldrich) in DMF (VWR). Commercially available N-Fmoc and side chain protected amino acids (5 eq, P3 Bio Systems) were activated for 5 min with HBTU (4.5 eq, ChemImpex) and DIEA (10 eq, Sigma) and then coupled to the resin for 1 hr. For beta-branched amino acids, 1.5 hrs was used. After successful coupling, the terminal Fmoc group was removed with 20% v/v piperidine in DMF for 15 min and then for an additional 15 min with fresh 20% piperidine in DMF. When peptides were completed, they were cleaved from the resin and side chains deprotected by a TFA cocktail (95:2.5:2.5 TFA/H2O/Triisopropylsilane) for 3 hrs rocking at room temperature. The peptide was then diluted ∼1/10 in cold diethyl ether and precipitated overnight (−80 °C). The resulting suspension was centrifuged (7,000 x g, 15 min, 4 °C) and the pellet was dried under compressed air. The pellet was then resuspended in 50:50 Buffer A:Buffer B, flash frozen, and lyophilized. This crude lyophilized material was purified by RP-HPLC over a C18 semi-preparative column (Phenomonex). Purified peptides were characterized by RP-HPLC (0-70% B gradient over 60 min) over an analytical C18 column (Higgins) and LC MS QTOF.

### *α*-Synuclein(gS87) synthesis

Lyophilized *α*-Synuclein C-terminal fragment **8** (5mM) and O-GlcNAc thioester peptide **7** (2eq) were dissolved in ligation buffer (200 mM NaH_2_PO_4_, 6M GnHCl, 25mM MPAA, 25 mM TCEP, pH 7.2) and allowed to react at room temperature overnight. Reaction progress was monitored by RP-HPLC. The reaction yielded pure O-GlcNAcylated *α*-Synuclein fragment **9** (85-140) after RP-HPLC purification. Product was confirmed by LC MS QTOF. The N-terminal thiazolidine (NThz) protecting group of **9** was removed with 200 mM methoxyamine pH 3.5 overnight to yield the free N-terminal cysteine fragment. Upon completion the reaction was purified by RP-HPLC and lyophilized. Product was confirmed by LC MS QTOF. Lyophilized **10** (9 mM final concentration) was dissolved in freshly prepared activation buffer (6M GnHCl, 200mM MPAA, 2.5eq acetylacetone). This solution was agitated at 200rpm for 3hrs at 25 °C. The reaction was monitored via RP-HPLC. Confirmation of the activated MPAA thioester was confirmed via LC-MS QTOF. Lyophilized O-GlcNAcylated *α*-Synuclein fragment **9** (85-140) (2eq) was dissolved in concentrated ligation buffer (6M GnHCl, 400mM NaH_2_PO_4_, 50mM TCEP) and added to the activation buffer containing thioester (1-84). The pH of the solution was adjusted back to 7.2 with concentrated NaOH. The reaction was complete after 24 hr at room temperature and fresh TCEP was added to reduce any disulfides before purification via RP-HPLC. After lyophilization, full length O-GlcNacylated *α*-Synuclein **11** (1-140) was confirmed via LC-MS QTOF. Radical desulfurization was used to convert the two cysteines in **11** to native alanines using VA-044. Protein **11** was dissolved in N_2_-sparged buffer (200mM NaH_2_PO_4_, 6M guanidine HCl, 175mM TCEP, pH 7.0). VA-044 and GSH (40mM and 80mM final concentration respectively) were dissolved to a final protein concentration of 2mM. The reaction was heated to 37 °C for 3 hrs and then purified by RP-HPLC to yield synthetic, full-length O-GlcNAc *α*-Synuclein [*α*-Synuclein (gS87) A69C F94W].

### *α*-Synuclein(gT72) synthesis

Expressed *α*-Synuclein N-term thioester **1** (1-68) (5 mM) and gT72 modified peptide fragment **2** (69-75) (2eq) were dissolved in ligation buffer and allowed to react at room temperature overnight at pH 7. Reaction progress was monitored by RP-HPLC and product formation was monitored via LC-MS QTOF. Product was purified by HPLC to yield pure O-GlcNAcylated *α*-Synuclein fragment **3** (1-75). Lyophilized **3** (9 mM) was dissolved in freshly prepared activation buffer and monitored for thioester formation. After thioester formation was confirmed via LC-MS QTOF, lyophilized *α*-Synuclein recombinant fragment **5** (76-140) (2eq) was dissolved in concentrated ligation buffer and the pH of the solution was adjusted back to 7.2 with concentrated NaOH. The reaction was purified by RP-HPLC to yield full-length *α*-Synuclein **6**. The radical desulfurization was carried out identically to the previously described method to yield synthetic, full-length O-GlcNAc *α*-Synuclein [*α*-Synuclein (gT72)].

### Aggregation Kinetic Experiments

ThT stocks were prepared by dissolving 1mM ThT in PBS and then sterile filtered through regenerated cellulose .22µm filters (VWR). Monomeric *α*-Synuclein was prepared by resuspending lyophilized protein in PBS buffer (Gen Clone (-) Ca, (-) Mg) and bath sonicated for 15 minutes, after which the solution was clarified by centrifugation at 20,000 x g for 30 minutes. The solution was then spin filtered through Microcon DNA fast flow Ultracel regenerated cellulose columns for 20 minutes to remove oligomers. The *α*-Synuclein monomer and ThT stock solution were combined to a concentration of 100 µM monomer and 10 µM ThT in PBS. 120µL of each solution was pipetted into the wells of a clear-bottomed black 96 well plate. Prior to assay use, the BioTek Cytation 5 instrument was preheated to 37 °C. The excitation wavelength was set to 450nm, and emission to 482nm for ThT fluorescence monitoring over the course of 168hrs with readings taken every 15 minutes with linear constant (1000rpm) shaking. Experiments were done in triplicate.

### Trp-Cys contact quenching measurements

All measurements were carried out at pH 7.4 and 37°C. Lyophilized powders of *α*-Syn, *α*-Syn(gT72) and *α*-Syn(gS87) were dissolved in 20mM Sodium Phosphate buffer, sonicated for 15 min and the solutions were centrifuged at 12000rpm for 2 min to get rid of the insoluble fractions. 20mM Sodium Phosphate buffer with and without 50% w/v sucrose were bubbled separately with N_2_O for an hour in sealed vials to deoxygenate and to scavenge solvated electrons that can be created by the UV laser pulse.

Decay data was collected using a pump-probe spectroscopy setup coupled with a syringe pump system (**Figure S2**) that injects protein samples into a sealed flow cuvette with specified concentrations of sucrose to vary viscosity (see **Table S1)** The Trp was excited using a 10ns UV pulsed laser at 289nm created from the fourth harmonic of an Nd:YAG laser (Continuum) and a 1-m Raman cell filled with 450 PSI of D_2_ gas. The lifetime of the Trp population was probed by transient absorption using a LASEVER 445 nm diode laser.

Sample viscosity was controlled using sucrose, which was varied using the syringe-pump system that mixed protein, buffer and a solution of buffer with 50% sucrose. A total volume of 1.2ml of the protein, the buffer and the sucrose solution were injected into a sealed cuvette using automated syringe pumps at a rate of 0.8ml/min and mixed for 3min. Measurements were taken for 0, 10, 20 and 30% w/v of sucrose at final concentrations of 20µM, 16µM and 12µM of *α*-Syn, *α*-Syn(gT72) and *α*-Syn(gS87) respectively. All measurements were obtained within ∼1hr of sample preparation and repeated twice. The viscosity of the injected volumes was confirmed using a BROOKFIELD DV-II+ Pro viscometer (**Table S1**).

## Acknowledgements

This work was supported by the National Science Foundation MCB 00581612 to L.J.L and the National Institutes of Health R01GM114537 to M.R.P.. qTOF analysis was performed at the USC Chemistry Mass Spectrometry Core and the Agilent Center of Excellence in Biomolecular Characterization.

## Notes

### Competing Interest Statement

The authors have declared no competing interest.

