## Supporting Information for "Post-translational Modification of α-Synuclein Modifies Monomer Dynamics and Aggregation Kinetics"

**Table of contents:**

**Figure S1.** Characterization of α-Syn proteins.	**Page S2**

**Figure S2.** Pump-probe spectroscopy coupled with a syringe pump system.	**Page S2**

**Table S1.** Measured viscosities of the samples for each sucrose percentage at 37 ℃.	**Page S3**

**Table S2.** Computed rates and diffusion coefficients of α-Syn, α-Syn(gT72) and

α-Syn(gS87) at 37 ℃.	**Page S3**

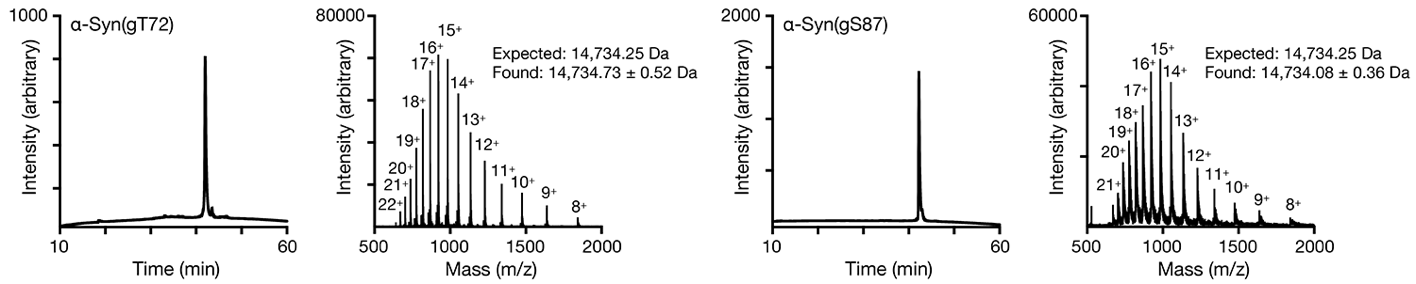

**Figure S1. Characterization of α-Syn proteins.** The indicated proteins were analyzed by RP-HPLC and ESI-MS.

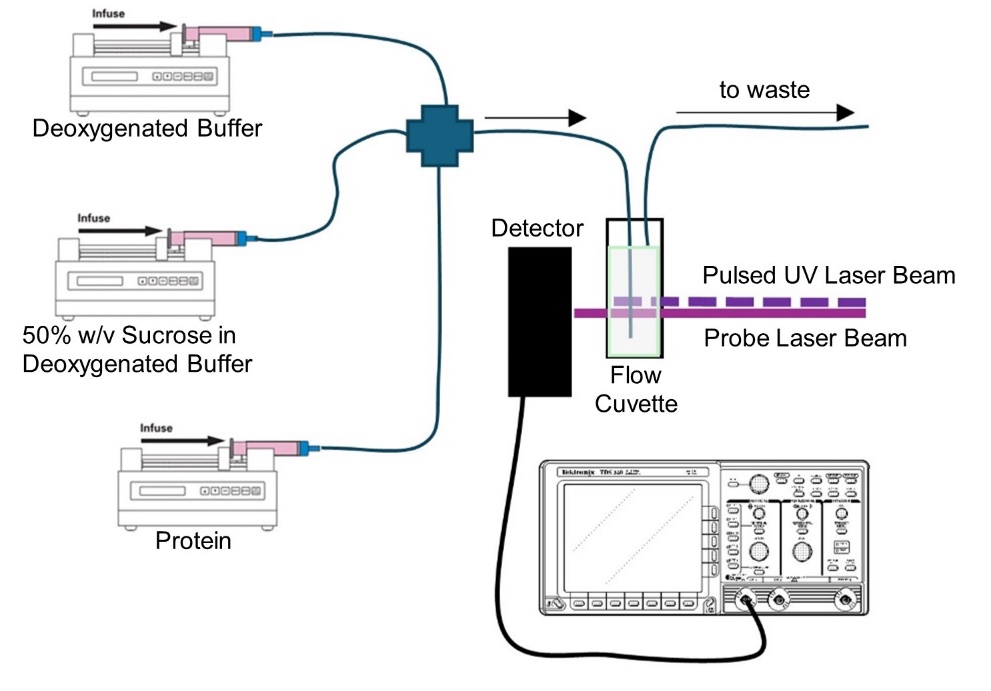

**Figure S2.** **Pump-probe spectroscopy coupled with a syringe pump system.**

To minimize the total sample usage, we used a syringe pump system with a sealed cuvette (800µL) for Trp-Cys measurements. Three syringe pumps (KDScientific-200) were used to inject the protein and the two deoxygenated buffers (0% and 50% sucrose) through a system of tubes where they mix before reaching the cuvette and further mixed while inside the cuvette by a stir bar for 3 min before measurement (see **Figure S2)**. To prevent sample contact with air, the solutions are loaded into gas tight syringes (Hamilton) and the cuvette is sealed. At the start of the measurements, air is flushed out by injecting 2ml of buffer. During data collection, a total volume of 1.2 mL was injected, consisting of 0.2 mL 300 μM protein, and a mixture of 0% and 50% sucrose solutions to create the sucrose concentrations listed in **Table S1.** Every measurement was taken on a freshly injected sample and injected at a rate of 0.8ml/min.

**Table S1.** Measured viscosities of the samples for each sucrose percentage at 37℃.

| **Sucrose %w/v** | **Viscosity (cP)** |
| --- | --- |
| 0 | 0.68 |
| 10 | 0.90 |
| 20 | 1.20 |
| 30 | 1.70 |

**Table S2.** Computed rates and diffusion coefficients of α-Syn, α-Syn(gT72) and α-Syn(gS87) at 37℃.

|  | **k_R_ (s^-1^)** | **k_D+_ (s^-1^)** | **D (cm^2^s^-1^)** |
| --- | --- | --- | --- |
| **α-Syn** | 1853891.85 ± 494561.52 | 2466415.38 ± 502962.75 | 4.21 ± 0.86 × 10^-7^ |
| **α-Syn(gT72)** | 1142347.16 ± 270348.86 | (LL) 2747859.19 | (LL) 7.78 × 10^-7^ |
| **α-Syn(gS87)** | (LL) 2118096.82 | 1084261.12 ± 109290.91 | (UL) 1.61 × 10^-7^ |

LL – Lower Limit, UL – Upper Limit
